# Kernel-based Nonlinear Manifold Learning for EEG Channel Selection with Application to Alzheimer’s Disease

**DOI:** 10.1101/2021.10.15.464451

**Authors:** Rajintha Gunawardena, Ptolemaios G. Sarrigiannis, Daniel J. Blackburn, Fei He

## Abstract

For the characterisation and diagnosis of neurological disorders, dynamical, causal and crossfrequency coupling analysis using the EEG has gained considerable attention. Due to high computational costs in implementing some of these methods, the selection of important EEG channels is crucial. The channel selection method should be able to accommodate non-linear and spatiotemporal interactions among EEG channels. In neuroscience, different measures of (dis)similarity are used to quantify functional connectivity between EEG channels. Brain regions functionally connected under one measure do not necessarily imply the same with another measure, as they could even be disconnected. Therefore, developing a generic measure of (dis)similarity is important in channel selection. In this paper, learning of spatial and temporal structures within the data is achieved by using kernel-based nonlinear manifold learning, where the positive semi-definite kernel is a generalisation of various (dis)similarity measures. We introduce a novel EEG channel selection method to determine which channel interrelationships are more important for the in-depth neural dynamical analysis, such as understanding the effect of neurodegeneration, e.g. Alzheimer’s disease (AD), on global and local brain dynamics. The proposed channel selection methodology uses kernel-based nonlinear manifold learning via Isomap and Gaussian Process Latent Variable Model (Isomap-GPLVM). The Isomap-GPLVM method is employed to learn the spatial and temporal local similarities and global dissimilarities present within the EEG data structures. The resulting kernel (dis)similarity matrix is used as a measure of synchrony, i.e. linear and nonlinear functional connectivity, between EEG channels. Based on this information, linear Support Vector Machine (SVM) classification with Monte-Carlo cross-validation is then used to determine the most important spatio-temporal channel inter-relationships that can well distinguish a group of patients from a cohort of age-matched healthy controls (HC). In this work, the analysis of EEG data from HC and patients with mild to moderate AD is presented as a case study. Considering all pairwise EEG channel combinations, our analysis shows that functional connectivity between bipolar channels within temporal, parietal and occipital regions can distinguish well between mild to moderate AD and HC groups. Furthermore, while only considering connectivity with respect to each EEG channel. Our results indicate that connectivity of EEG channels along the fronto-parietal with other channels are important in diagnosing mild to moderate AD.

## I. Introduction

The electroencephalogram (EEG), even though recorded at the scalp level, reflects the grossly summed currents of the electrical fields generated by the neural activity in the cortical neural circuits. Through the EEG, the behaviour and integrity of underlying neural circuits can thus be indirectly studied [1]. The analysis of hidden structures within EEG data is therefore important and has gained considerable attention [2].

Numerous EEG studies have previously revealed the importance of nonlinear methods for the diagnosis of neurological disorders [2]. Among these, in-depth dynamical analysis, such as the analysis of linear and nonlinear dynamic inter-relationships between EEG channels, causality and cross-frequency coupling analysis, has gained considerable attention [2–6]. However, some of these methods can often incur high computational costs. Consequently, in practice, the selection of important EEG channels is vital [7] before conducting an in-depth dynamical analysis of high dimensional EEG data. Furthermore, to select channels to conduct non-linear dynamical analysis, the channel selection method should be able to account for non-linear interactions between the channels.

In EEG channel selection techniques, usually features from the channels are extracted and important channels are selected via feature selection. These feature selection methods can be categorised into the following three groups [7]. a) *Filtering methods*: Independent evaluation criteria, such as distance measures, entropy, mutual information, are used for the purpose of channel selection. Depending on the criteria used, these are often only based on either single or pairwise EEG channel(s). Filtering methods are good at eliminating irrelevant and redundant features. b) *Wrapper methods*: Subsets of features are generated based on a method of choice. Each subset is evaluated using a classification algorithm to select a subset of channels. These are based on greedy search algorithms aiming to find the best possible combination of features. c) *Embedded Methods*: These techniques do the feature selection and the classification simultaneously. For example, LASSO based feature selection, logistic regression and decision-tree are techniques that come under embedded methods. Ranking of features can be easily done using embedded methods.

The brain is a complex nonlinear system, and the nonlinear nature of brain dynamics can be readily detected considering the inter-relationships between channels [2]. On the other hand, the analysis of nonlinear structures locally and globally within EEG data can also be used for channel selection purposes [8], which is often implicitly considered in many EEG channel selection approaches discussed above. In this study, a novel EEG channel selection method based on spatio-temporal linear and nonlinear EEG inter-relationships is presented. This is achieved by using pairwise representation [9] via kernel-based manifold learning (dimensionality reduction). The features based on spatio-temporal pairwise distance measures are statistically compared to choose a subset of features that are significantly different between AD and HC groups. This subset is then assessed using linear SVM classification with Monte-Carlo cross-validation to evaluate which features are better at differentiating between AD patients and HCs. Furthermore, the linear SVM classification weights are used to rank the selected pairwise features. Therefore, the method presented is a hybrid form [7] of the aforementioned three categories of feature selection methods.

Pairwise representation has gained attention in various fields, such as bioinformatics, neuroscience, cognitive psychology, social sciences, text and web mining [10–15], as it can be used to model highly nonlinear data and to reveal many hidden structures within the data [9]. In pairwise representation, various distance measures relating to the concepts of (dis)similarity within the data are considered. Similarity or dissimilarity between two features or variables, in general, express the degree to which the two objects are respectively alike/related or different/distinct. Cross-correlation, Euclidean and Manhattan distances are examples of (dis)similarity measures. These can be used for the pairwise comparison of objects in the space of measured variables [16]. Therefore, (dis)similarity can be used as a measure of synchrony and linear/nonlinear inter-relationships [15, 17]. In neuroscience, brain functional connectivity (synchrony) is characterised using different measures of (dis)similarity [15, 18–25]. Thus, regardless of structural connectivity, brain regions functionally connected under one measure do not necessarily imply the same with another measure as they could even be disconnected [8, 15]. Dauwels *et al*. [8] showed that various (dis)similarity measures can be correlated to each other, such as in the application of early diagnosis of AD. Hence, these correlated measures can often be grouped, and one measure from each group is sufficient to analyse the structures within the data [8]. In this study, a spatio-temporal pairwise representation is applied using kernel-based nonlinear manifold learning and the focus is on functional connectivity changes (synchrony) and thereby EEG channel selection. The motivation behind utilising a kernel-based method is based on the fact that positive semi-definite (PSD) kernels are a generalisation of distance measures [26]. In this work, the kernel matrix is evaluated using nonlinear manifold learning via Gaussian Process Latent Variable Model (GPLVM) [27]. Robust kernel Isomap [28] is used as an initialisation method for GPLVM (Isomap-GPLVM). This enables the learning of both local similarities and global dissimilarities within the EEG data and embedding it in the reduced-dimension manifold (latent space) [29]. Furthermore, since dimensionality reduction is used to reduce the temporal dimension, temporal structures within the data are taken into account. This is not possible in the direct use of popular distance measures, as these methods only consider the spatial aspect of the data and not the temporal aspect unless a delay embedding is used.

In summary, the kernel matrix evaluated, using Isomap-GPLVM is PSD and can be regarded as a more objective (dis)similarity measure containing information on both linear and nonlinear spatio-temporal EEG interrelationships. Therefore, a generalisation of different functional connectivity measures [26], and can be a better alternative to using various (dis)similarity measures each from the respective groups as mentioned earlier. Since statistical comparisons of kernel matrices between groups are used to select a subset of functional connectivity features, which are then evaluated using linear SVM, the proposed approach is an integration of filtering and wrapper methods. It is shown that the channels pairs chosen from our approach can be linked to other EEG studies in the literature considering connectivity analysis in AD.

This paper is organised as follows. Subsection I A provides a short list of definitions for multi-disciplinary concepts used in this paper. Section II provides specifics of the EEG data used and the pre-processing steps. Section III discusses the manifold learning methodology via Isomap and GPLVM and the use of related kernel-based (dis)similarity matrix for the classification of EEG data, which are measured from a group of AD patients and an age-matched healthy control cohort. This section also presents the statistical comparison and the correction method used to deal with the multiple hypothesis testing problem. Linear SVM and Monte-Carlo cross-validation procedures are presented. Section IV discusses the results obtained followed by the concluding remarks and the future work in Section V.

### A. Definitions of Concepts

#### (Dis)similarity

Similarity or dissimilarity between two EEG channels expresses the degree to which the two are respectively related or distinct [10]. Measures of statistical dependency, such as crosscorrelation, mutual information and geometric distance measures fall under (dis)similarity.

#### Functional brain connectivity

Statistical dependence between EEG channels [30]. Quantified using various (dis)similarity measures. Different measures may not necessarily mean the same [15].

#### Effective connectivity

How one electrode affects the EEG rhythms at another electrode, as a reflection of causal interaction between the two underlying cortical regions [30].

#### Synchrony

Functional and effective interactions between EEG channels. For example, Crossfrequency interactions between different frequency ranges across EEG channels [2].

#### Complexity

Characterise connections between EEG channels. Taken from network theory [31]. (Dis)similarity between EEG channels can be used as one measure to characterise complexity.

#### Single channel complexity (signal complexity)

Linear and nonlinear information contained within an EEG channel [32]. For example, entropy, auto-mutual information [2] and cross-frequency interactions [33] within the signal (e.g. within-frequency amplitude coupling).

## II. Data

In this work, we include a total of 20 AD cases and 20 age and gender-matched healthy controls (HC) (less than 70 years of age) all selected based on clinical and radiological diagnostic criteria as described in [34]. Task-free EEG recordings that require minimal cooperation of AD patients are used; typically, this group of patients can have difficulty engaging with and following cognitive tasks. The details of experimental design, diagnosis confirmation, data acquisition and EEG electrode configuration are provided in [34]. All AD participants were in the mild to moderate stage of the disease at the time of EEG recordings.

The Sheffield Teaching Hospital memory clinic team, which focuses mainly on young-onset memory disorder, recruited all AD participants. AD participants were diagnosed between 1 month and 2 years before data collection. The diagnosis was conducted using a series of psychological tests, medical history, neuro-radiological examinations and neurological examinations. High resolution structural magnetic resonance imaging (MRI) was used to eliminate other causes of dementia in all participants. The age and gender-matched HC participants were recruited whose structural MRI scans and neuropsychological tests were normal. This study was approved by the Yorkshire and The Humber (Leeds West) Research Ethics Committee (reference number 14/YH/1070). All participants gave their informed written consent.

### A. EEG Data

A modified 10-10 overlapping a 10-20 international system of electrode placement method was utilised to acquire the EEG recordings. All EEG data were recorded using an XLTEK 128-channel headbox with Ag/AgCL electrodes placed on the scalp at a sampling frequency of 2kHz. A common referential montage with linked earlobe reference was used. During the 30 minutes of EEG recording, the participants were encouraged not to think about anything specific. All participants had their eyes-open (EO) for 2 minutes and then eyes-closed (EC) for 2 minutes, in repeat, during the 30-minute recording. EEG data were reviewed by an experienced neurophysiologist on the XLTEK review station with time-locked video recordings (Optima Medical LTD). Subsequently, from the resting-state EEG recordings, three 12-second artefact-free epochs under EO and EC conditions were isolated.

From the referential montage, the following 23 bipolar channels were produced for the analysis: F8–F4, F7–F3, F4–C4, F3–C3, F4–FZ, FZ–CZ, F3–FZ, T4–C4, T3–C3, C4–CZ, C3–CZ, CZ–PZ, C4–P4, C3–P3, T4–T6, T3–T5, P4–PZ, P3–PZ, T6–O2, T5–O1, P4–O2, P3–O1 and O2–O1. The bipolar channels were obtained by simply subtracting the two common referenced signals involved. To avoid any confusion, any bipolar channel(s) will be referred to from here on as *EEG channel(s)* or just *channel(s)* unless otherwise specified. In summary, three 12-second epochs of EO EEGs are collected from 20 HC and 20 AD participants and used in this study.

### B. Pre-processing Tasks

In this study, since the high-dimensional temporal structures of the multi-channel EEG are examined, the use of filters would pose a major issue due to the phase-related distortions induced [35]. Therefore, we first convert the time-series EEG data to the frequency-domain using the Fast Fourier Transform (FFT), unwanted frequency components can be easily removed with minimal phase distortions. Thereafter, the inverse-FFT was used to reconstruct the time-domain signals without the unwanted frequency components. The analysis in this work was conducted using EEG frequencies between 2 to 100Hz. Furthermore, frequency components around 50Hz (49.5-50.5Hz) were also removed to avoid any contamination by AC power line noise. After removing the unwanted frequency components, the reconstructed time-domain signals were then down-sampled to 200Hz.

## III. Methods

This paper presents a methodology where (dis)similarity information within EEG data via kernel-based manifold learning is used to determine the important EEG channel inter-relationships (channel pairs), in the case of AD. These channel pairs can be used for further analysis of cross-frequency coupling and nonlinear dynamical changes in the EEG between HC and AD groups.

Manifold learning is an approach to perform nonlinear dimensionality reduction where a lower-dimensional representation is learned from higher-dimensional data. The EEG is multi-channel time-series data (i.e. highdimensional spatio-temporal data). Kernel-based manifold learning can be used to reduce the temporal dimension, and determine both linear and nonlinear spatial and temporal structures within the EEG data. The kernel matrix obtained from such manifold learning methods will contain the (dis)similarity information within the original data, and named a kernel (dis)similarity matrix. This matrix can be used as a general measure of spatiotemporal functional connectivity–a measure of synchrony between EEG channels.

As mentioned in Section I, various measures of (dis)similarity, such as correlation and manhattan distance, can be used to analyse functional connectivity. While distance measures only look into the spatial structures within time-series data, (dis)similarity via kernel-based manifold learning can learn both spatial and temporal structures simultaneously.

Manifold learning methods that preserve local similarities within the data in the lower-dimensional space (or latent space) imply a mapping from the data space to the latent space [29]. Kernel principal component analysis (KPCA), locally linear embedding (LLE), t-SNE and Isomap are such methods. Conversely, methods that imply a mapping from the latent space to the data space preserve global dissimilarities, i.e. two points that are relatively distant in the data space are guaranteed to be placed at a distance in the latent space [29]. GPLVM is a kernel-based global dissimilarity preserving mapping. The kernel in kernel-based manifold learning methods, such as Isomap and GPLVM, describes the nonlinear structures within the data in a non-parametric fashion.

The methodology proposed in this study integrates the strength of both categories by employing GPLVM and Isomap, where Isomap is used as an initialisation method for GPLVM; we name it Isomap-GPLVM. As a result, the spatio-temporal local similarities and global dissimilarities within the EEG data are preserved in the latent space. The resulting kernel matrix from GPLVM will contain such information as a measure of (dis)similarity between EEG channels. Participant-specific kernel (dis)similarity matrices of HC and AD groups are statistically compared to determine which (dis)similarities between brain regions (EEG channels) are significantly different between the groups. Linear SVM is then used to determine the rank of the important (pairwise) EEG channel inter-relationships.

### A. Gaussian Process Latent Variable Model (GPLVM)

As a nonlinear manifold learning technique, a GPLVM [27] learns the mapping of a high-dimensional observed dataset **Y** *∈* ℝ^*N ×D*^ from the corresponding low dimensional latent space **X** *∈* ℝ^*N ×Q*^, *Q < D*, using a Gaussian Process (GP) [36], i.e. a mapping from **X** *→* **Y**. Here **Y** = [**y**_1_, *· · ·*, **y**_*N*_]^*T*^, **y**_*i*_ *∈* R^1*×D*^ and **X** = [**x**_1_, *· · ·*, **x**_*N*_]^*T*^, **x**_*i*_ *∈* ℝ^1*×Q*^. The GP mapping **X** *→* **Y** is governed by a kernel matrix **K**, where **Y** = *𝒢𝒫* (**X, K**). The likelihood of the data given the latent positions is 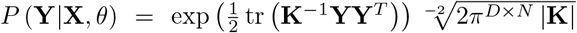 where **K** *∈* ℝ^*N ×N*^,

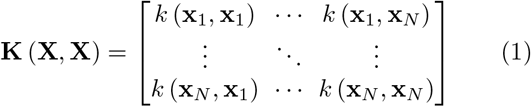

is a positive semi-definite matrix, where *k* (**x**_*i*_, **x**_*j*_) is the kernel/covariance function with a set of hyperparameters *θ*. The use of a kernel function allows the nonlinear functional mapping from **X** to **Y** and provides a probabilistic nonlinear latent variable model. In GPLVM, the maximising of the marginal log-likelihood, log *P* (**Y** | **X**, *θ*), is done with respect to both **X** and *θ*, hence the optimal estimates 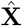 and 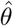 are obtained jointly. It should be noted that when using the Radial Basis Function (RBF) kernel with GPLVM, the length-scale hyper-parameter influence how much similarity or dissimilarity information from the data space is embedded within the latent space [29].

In order for GPLVM to optimise well and perform to a given criteria, appropriate initialisation of 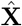 and initial conditions for 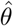 should be provided. As mentioned earlier, initialisation of 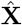 is done using Isomap, this preserves both local and global (dis)similarities of the spatiotemporal manifold of the EEG data. Initial conditions for the hyper-parameters 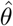 are chosen (via grid search) depending on how well HC and AD kernel (dis)similarity matrices are statistically distinguishable. This will be elaborated further in the following subsections.

### B. Isomap as an initialisation for GPLVM

Isomap [37] aims to preserve the geometry within nonlinear data by using the geodesic distances (along the surface of the high dimensional manifold) between the data points. It approximates the geodesic distances using weighted neighbourhood graphs to project the highdimensional data to a lower-dimensional manifold, preserving shape information [37]. This is the reason for the choice of Isomap over methods, such as KPCA and tSNE, as the initialisation for GPLVM. In this study, the robust kernel Isomap [28] variant is used.

Robust kernel Isomap approximates the geodesic distance to project the data into the latent space (i.e. lower-dimensional manifold), while preserving topological stability and providing a method for eliminating critical outliers [28]. The data points are projected according to how close the points are along the high dimensional manifold surface (i.e. local similarities). In the analysis of EEG data, robustness to noise is vital as this could affect the local similarities and the geodesic distance calculations. This is the main reason for utilizing robust kernel Isomap instead of the competing LLE method and its variants.

### C. Kernel-Based nonlinear Manifold Learning of High-dimensional EEG Data Using Isomap-GPLVM

The GPLVM is a global dissimilarity preserving mapping [29]. To also preserve the local similarities, we employ another manifold learning technique, the robust kernel Isomap [28], which is used as an initialisation for the estimated latent space 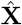 in the GPLVM optimisations (see Section III A). In such a way, both local similarities and global dissimilarities are preserved within the latent space. To demonstrate this, Fig. 1 illustrates Isomap-GPLVM operating on a *coarsely sampled* (i.e. dataset has only fewer samples) swiss-roll dataset.

**FIG. 1.**
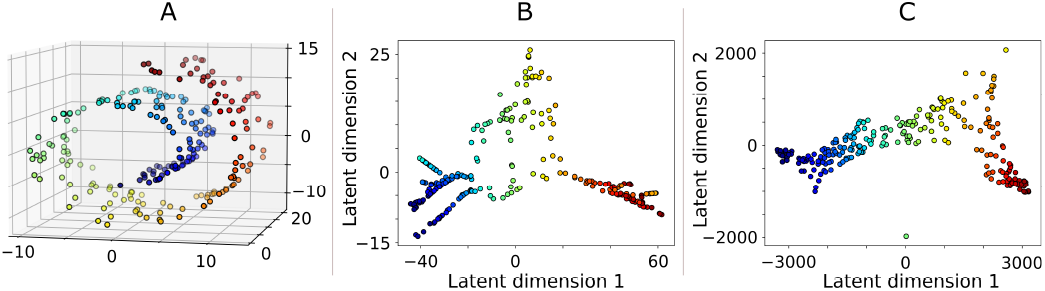
Isomap, used as an initialisation method for GPLVM, can preserve both local similarities and global dissimilarities, of the data, within the latent space. **A)**. The original Swiss Roll dataset. Data coarsely sampled from the original high dimensional manifold. **B)**. Isomap projection preserves local similarities in the data structure from *Data space* to *Latent space* mapping. **C)**. GPLVM optimises the lower-dimensional manifold produced by Isomap, where global dissimilarities in the data structure are embedded into the latent space through a *Latent space* to *Data space* mapping.

Robust kernel Isomap approximates the geodesic distance to project the data into a lower-dimensional manifold preserving local similarities, as shown in Fig. 1B. GPLVM uses the Isomap projection of the data as an initialisation of the latent space. GPLVM then operates on this latent space and optimises it according to the distance between data points, along the high-dimensional manifold surface (i.e. global dissimilarities), in the original data space (Fig. 1C). The kernel matrix **K** (**X, X**), from the optimised GPLVM, governs the latent space to data space mapping (see Section III A). This kernel matrix contains both the local and global (dis)similarity information within the original data – kernel (dis)similarity matrix.

Isomap-GPLVM is used to analyse the EEG data by reducing the temporal dimension. The manifold learning methods used will learn the local and global (dis)similarities (Isomap–local similarities and GPLVM– global dissimilarities) along the spatio-temporal manifold of the EEG data. Therefore, the kernel matrix, **K** (**X, X**) *∈* ℝ^23*×*23^, contains the spatio-temporal (dis)similarity information between the 23 respective EEG channels.

Isomap-GPLVM is applied individually to the EEG data of each AD and HC participant, obtaining a kernel (dis)similarity matrix for each participant. Following the definition of the data space **Y** *∈* ℝ^*N ×D*^ (as given in Section III A), here *N* = 23 (23 EEG channels, Section II A) and *D* is the temporal dimension that is to be reduced. The RBF was used as the kernel function. The respective latent space of each AD and HC participant will be **X** *∈* ℝ^*N ×Q*^. There are two reasons for setting the latent dimension *Q* = 7: i) to achieve a nearly 100% recovery accuracy in the GPLVM mapping to the original data space (i.e. from **X** to **Y**); and ii) to achieve the best results in distinguishing between AD and HC. Before applying nonlinear manifold learning, each EEG channel of all participants was de-meaned and normalised, such that the absolute maximum value attained by the respective channels is 1.

It should be noted that the pre-processed 23-channel EEG data of each participant contains 2400 time samples (see Section II A), **Y** *∈* ℝ^23*×*2400^. This can be nearly perfectly represented in the latent space with only 7 latent variables using Isomap-GPLVM, **X** *∈* ℝ^23*×*7^. To achieve the same recovery accuracy, linear latent representation method, such as principal component analysis, requires 20 latent variables.

### D. Statistical Comparison of Kernel (dis)similarity matrices

Using kernel (dis)similarity measures, cortical regions (EEG channels) that have higher *synchrony changes* between patient and control groups are determined. A kernel (dis)similarity matrix for each HC and AD participant is evaluated using Isomap-GPLVM.

Kernel (dis)similarity matrices from the HC group are statistically compared against the AD group to understand the global functional connectivity differences between the groups. This is achieved through element-wise comparison of matrices from the two groups. The elements of all the kernel (dis)similarity matrices are compared between AD and HC groups. This is done to determine if there is a significant difference (i.e. *p*-value *<* 0.05) between the groups with respect to each element. If so, the corresponding elements in the significance matrix, **S** *∈* ℝ^23*×*23^, are set to 1, indicating that these channel pairs have a statistically significant difference in (dis)similarities (inter-relationships) between HC and AD groups, and 0 otherwise. Therefore, **S** indicates (at an EEG sensor level) that underlying cortical global functional connectivities vary significantly between HC and AD. This information can be used for selecting pairs of channels for model-based signal processing and analysis.

The Mann–Whitney U test [38] is used for statistical comparison–as the distribution of individual elements of the kernel matrices within either group is non-Gaussian. Due to the multiple statistical comparisons used here, the *p*-values need to be approximately corrected. Also, due to the high number of comparisons (23 EEG channels, 253 channel combinations), controlling of the false discovery rate (i.e. positive results that could be in fact negative) [39]) is preferred over family-wise error rate controlling [8, 40]. The Benjamini-Hochberg [39] false discovery rate controlling (FDR) method is used to adjust the *p*-values to obtain a corrected significance matrix, 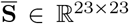 ∈ ℝ^23*×*23^.

### E. GPLVM Kernel hyper-parameter initial condition determination

The (dis)similarity measures from the kernel matrix of GPLVM depend considerably on the initial conditions of the RBF kernel hyper-parameters *θ*. The initial conditions are determined using a grid search method using the corrected significance matrix 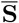 (Section III D).

The best set of initial conditions for *θ* are chosen based on how well the kernel (dis)similarity matrices are distinguishable from HC to AD (i.e. the highest number of corrected significant *p*-values, 1’s in 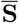). This was done for several latent dimensions, i.e. *Q* = [5, 8] and *Q* = 7 was chosen. The length-scale hyper-parameter (when using RBF kernel in GPLVM) influences how much similarity and dissimilarities information is embedded in the latent space (Section III A). Therefore, the initial conditions determined, as mentioned above, leads the manifold learning method to produce a kernel matrix where its (dis)similarity measures are relatively specific for the differentiation of HC and AD EEG data.

It is important that 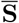 rather than **S** is used for initial condition determination. The Benjamini-Hochberg method is conservative (in correcting *p*-values) relative to other FDR methods[41]. Therefore, initial conditions determined using 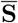 (highest number of 1’s in 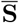) ensures that GPLVM produces kernel matrices that are statistically well distinguishable between the two groups.

### F. Kernel (dis)similarity matrix based global functional connectivity changes and channel pair selections

The pairwise (dis)similarities from the kernel matrices are used as features to determine which EEG channel pairs are more *important globally* (considering interconnections with all the other channel pairs) in differentiating between HC and AD groups. These features are processed using linear SVM classification and the weights of the linear SVM classifier are used as a measure of the *global importance* of a channel pair. There are 23 EEG channels (Section II A), therefore a total of 253 channel combinations – 253 (dis)similarity features. These features can be easily obtained from either the lower or upper triangular elements in the kernel (dis)similarity matrices as shown in Fig. 2A.

**FIG. 2.**
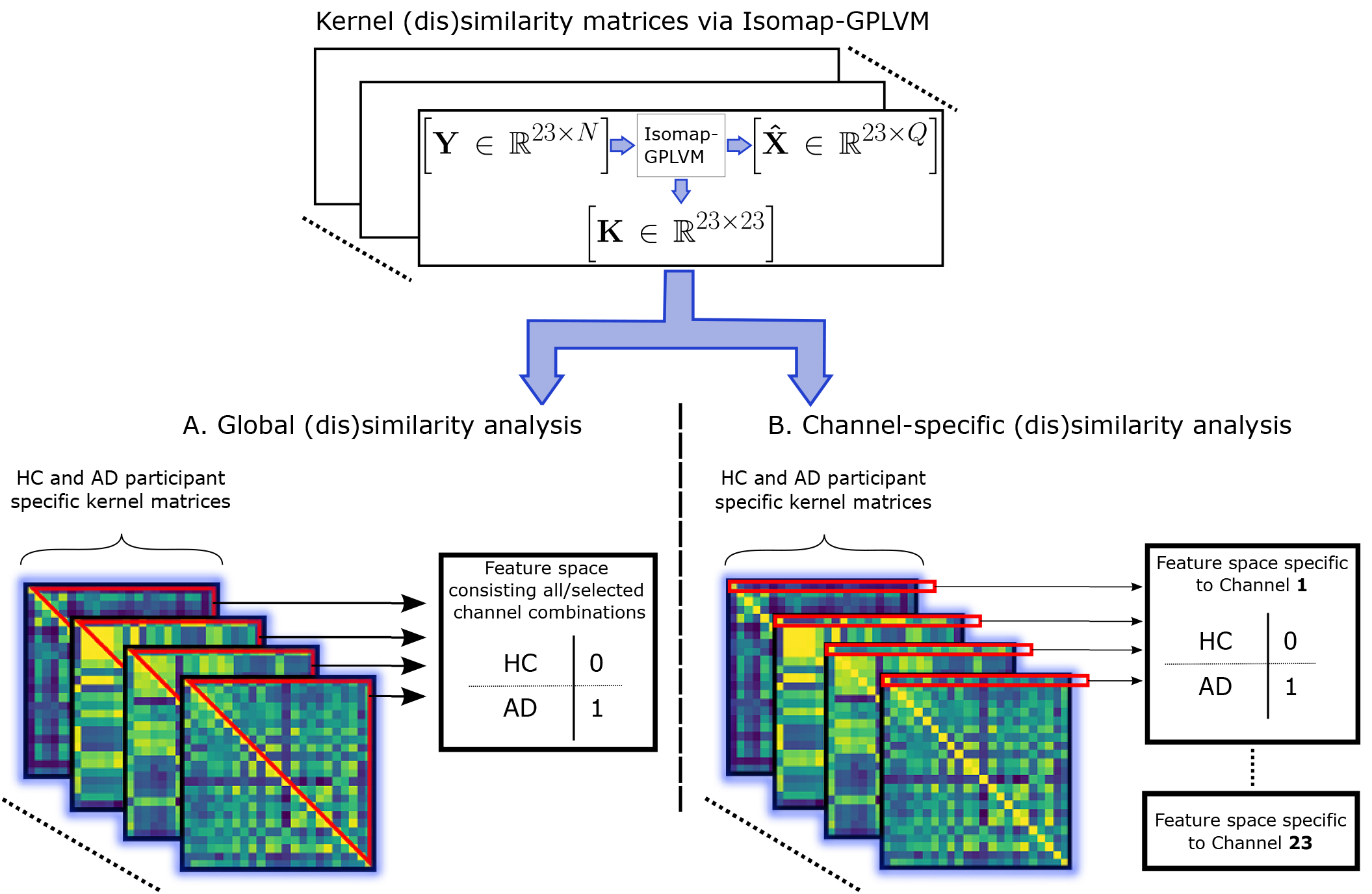
(Dis)similarity analysis via kernel based manifold learning– methodology. Participant specific kernel (dis)similarity matrices are evaluated using Isomap-GPLVM. From the EEG data, **Y**, Isomap-GPLVM learns the spatiotemporal local similarities and global dissimilarities within the data (see Section III C). This information is embedded in the resulting latent space 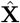 and is reflected in the kernel matrix **K** (see Section III A). **K** governs the GPLVM mapping 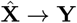. (Dis)similarity analysis using **K** is conducted in two approaches. **A). Global (dis)similarity analysis methodology of AD and HC EEG data**. From all channel pair combinations, selected pairs using **S** are used as features. The feature space has two classes, AD and HC. The classification is binary – AD is denoted as 1 and HC as 0. A linear SVM classifier is used on the feature space to determine which channel pairs (inter-relationships) are better at distinguishing between groups in a global sense. **B). Channel-specific (dis)similarity analysis methodology of AD and HC EEG data**. All channel-specific rows from these matrices are grouped into channel-specific feature spaces. Each feature space has two classes, AD and HC. Individual linear SVM classifiers are used on each feature space to determine which channels and their respective inter-relationships with other channels are better at distinguishing between the groups.

The linear SVM classification can be carried out under several different settings:

1. Using all 253 channel pair combinations.
2. Using only a subset of channel pair features from either **S** or 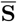.
3. Using a L1 or L2 penalty under either setting 1) or 2).

The area under the receiver operating characteristic curve (AU-ROC) is used as a performance metric to compare the above procedures. The subset of features selected using **S** along with a L2 penalty results in the best AU-ROC value. The AU-ROC is obtained using a Monte-Carlo cross-validation scheme (elaborated in Section III H) for the linear SVM classifications.

This analysis is used to determine the *global importance* of channel pairs. Considering all available EEG channels, which spatio-temporal inter-relationships between different cortical regions (EEG channels) are more important in differentiating between HC and AD groups.

### G. Kernel (dis)similarity matrix based channel-specific functional connectivity changes and channel pair selections

In the previous subsection, a global approach to determining the important inter-relationships in the EEG was presented. However, it is also important to understand how a particular underlying region of the cortex interacts with other regions, i.e. if there are any significant region-specific synchrony changes between HC and AD groups [42–44]. Therefore, in this subsection, a channel-specific approach is presented. More specifically, considering a particular channel and only its inter-relationships with other channels, to determine which channel-specific connectivities can distinguish well between AD and HC groups. This provides another layer of information in determining important functional connectivity changes in AD.

Each row in the kernel matrix contains a measure of (dis)similarity of a particular channel against all other respective channels. Thus, channel-specific rows from all the participant-specific kernel (dis)similarity matrices are grouped into 23 different feature spaces (23 EEG channels, Section II A). Each channel-specific feature space will contain 22 features, i.e. (dis)similarity measures against other respective channels. These feature spaces are assessed individually using linear SVM. Therefore, 23 different linear SVM classifiers are used to make a channel-specific assessment to determine how each channel and the respective inter-relationships (channel-specific pairs) change in AD. Fig. 2B illustrates this methodology.

The AU-ROC of each feature space is used to determine which channel-specific feature spaces can better distinguish between HC and AD groups. Similar to Section III F, it was found that the subset of channel-specific features selected using an L2 penalty resulted in the best AU-ROC values across all feature spaces. The application of linear SVM to channel-specific feature spaces gives a channel-specific assessment of important changes in spatio-temporal channel synchrony between HC and AD. The normalised average linear SVM weights of each channel-specific feature space give a ranking to the synchrony changes with respect to the channel.

### H. Linear SVM and Monte-Carlo cross-validation procedure

As mentioned in Section II A, this study comprises of 20 HC and 20 AD participants. From each participant, three 12-seconds epochs of EO EEGs are used. Kernel (dis)similarity matrices of the EEG data are produced for each AD and HC participant using Isomap-GPLVM, for all three epochs.

Monte Carlo cross-validation was used where, from the *first epoch*, 10 HC and 10 AD participants are randomly picked for the training set. The remaining 10 HC and 10 AD participants from the *first epoch* were used for testing. Since the brain is a highly complex and stochastic system [45], each epoch would have certain differences. Therefore, the testing dataset also included the 2^nd^ and 3^rd^ epochs of all participants. 1000 such random samples were made and the averaged AU-ROC from the *testing set* was used as the metric to determine the performance of the linear SVM classification. The average of the linear SVM weights from the 1000 samples are taken. The averaged weights are normalised such that the highest absolute averaged weight is 1. This is used to rank the features in the respective feature space. This procedure of linear SVM classification with Monte-Carlo cross-validation is used in sections III F and III G.

The use of the Monte-Carlo cross-validation strategy is because some AD participants could easily be detected while others maybe not. Since such information is not available, a randomised cross-validation strategy is necessary for obtaining a fair balance in linear SVM weights. Furthermore, the use of other epochs only in the testing set ensures the generalising capability of the linear SVM classifier and the weightings are skewed towards the most significant predictors.

It should be re-emphasised that the focus of this work is on channel selection. Linear SVM classification is only used to determine which channel inter-relationships exhibit the highest *synchrony changes* between patient and control groups. This is achieved using spatio-temporal (dis)similarities between channels as a measure of synchrony. The proposed methodology can also be further developed as a diagnostic tool for the detection of AD, provided a sufficiently large number of participants is available.

### I. Summary of Isomap-GPLVM and subsequent procedures

The Isomap-GPLVM method and the procedures for analysing the resulting kernel (dis)similarity matrices using linear SVM with Monte-Carlo cross-validation (SVMMCV) are summarised as follows:

1. Pre-process the data (Section II B).
  a. Apply Fourier transform method of filtering to remove unwanted frequency components.
  b. Normalised data such that the maximum amplitude of the signals are 1.
  c. Use Isomap as an initialisation for the latent space (Section III B).
2. Use the grid search method to determine the best kernel hyper-parameter, *θ*, initial conditions for GPLVM (Section III E).
  a. For a set of initial *θ*, GPLVM (with the Isomap initialisation of the latent space) is used to evaluate the kernel matrices for all participant data.
  b. All the HC kernel matrices are statistically compared against all the AD kernel matrices to obtain the **S** matrix.
  c. The 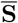 matrix is obtained by applying the Benjamini-Hochberg false discovery rate controlling method to account for the multiple statistical comparisons problem.
  d. The best initial condition is the one which results in the most dense 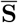. i.e. matrix with the most 1s indicating that those set of *θ* initial conditions produce kernel (dis)similarity matrices that are statistically well distinguishable between HC and AD.
3. Steps 2. and 3. above constitutes the IsomapGPLVM method.
4. SVM-MCV (Section III H) is used to analyse the kernel (dis)similarity matrices to identify the best channel-pairs that can differentiate between HC and AD, i.e. ranking the subset of pairwise channel (dis)similarities that have significant statistical differences between the two groups (**S**–step 3b). The channel-pair rankings are done using the linear SVM weights. Two approaches are used in applying SVM-MCV:
  a. Global analysis (Section III F). All pairwise (dis)similarities that are statistically significant are used. This identifies the best channel pairwise comparisons that can distinguish between HC and AD considering the whole EEG.
  b. Channel-specific (Section III G). Each row of all the kernel matrices are used to form a channel-specific feature space. SVM-MCV is applied to each feature space individually. This ranks the statistically significant channel pairs considering a specific channel and its relations with the rest of the EEG.

## IV. Results and Discussion

The Isomap-GPLVM method introduced in Section III was applied to the EO EEG data (Section II A). The best set of initial conditions for the hyper-parameters, *θ*, was determined by using the three 12-second epochs via the procedure explained in Section III E, using the corrected significance matrix 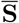 (Section III D). From the kernel (dis)similarity matrices evaluated, the channel interrelationships that were able to differentiate well between AD and HC groups are presented in this section. This was done by using two different kernel (dis)similarity matrix based approaches via linear SVM. The first is based on global functional connectivity changes (as described in Section III F) and the second is based on channel-specific functional connectivity changes (as described in Section III G).

This study is conducted using 20 HC and 20 AD participants. Each participant has three sets of 12-second epochs, except for one AD participant with only one epoch from this participant available. Fig. 3 illustrates the bipolar montage EEG channels used in this work in a 10-20 international standard electrode placement map. Table I shows the 23 bipolar montage EEG channels used and the respective underlying cortical regions.

**FIG. 3.**
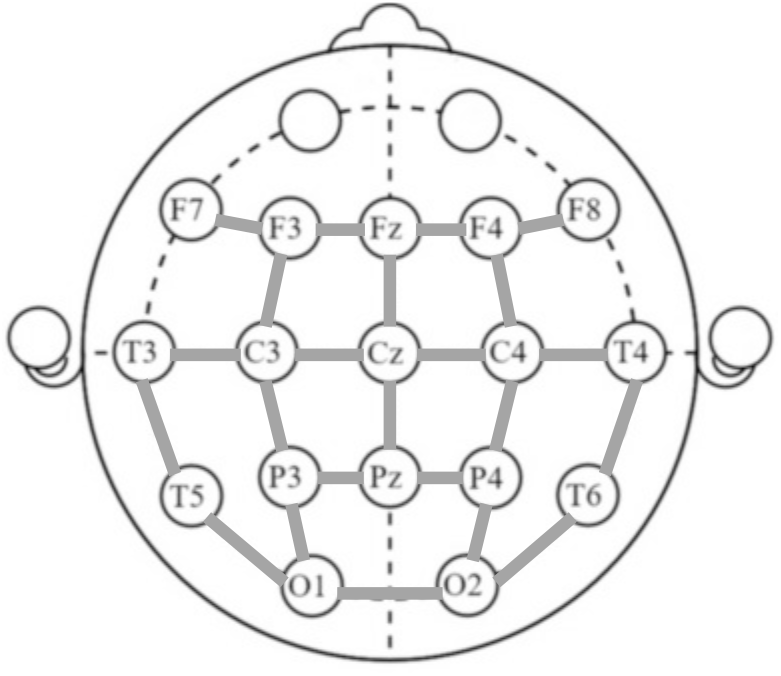
All the 23 channels, bipolar montage. EEG bipolar montage channels mapped into a 10-20 international standard arrangement. The bold grey lines connecting any two EEG nodes indicate that these two nodes result in a bipolar channel.

**TABLE I.** List of all 23 channels of the scalp EEG bipolar montage and the corresponding underlying cortical regions

| Channel index and name | Corresponding cortical region |
| --- | --- |
| 0 O1-O2 | Occipital |
| 1 P4-O2 |  |
| 2 P3-O1 |  |
| 3 T5-O1 |  |
| 4 T6-O2 |  |
| 5 P3-PZ | Parietal |
| 6 P4-PZ |  |
| 7 T3-T5 | Temporal |
| 8 T4-T6 |  |
| 9 C3-P3 | Centro-Parietal |
| 10 C4-P4 |  |
| 11 CZ-PZ |  |
| 12 C3-CZ |  |
| 13 C4-CZ |  |
| 14 T3-C3 | Centro-Temporal |
| 15 T4-C4 |  |
| 16 F3-C3 | Fronto-Central |
| 17 FZ-CZ |  |
| 18 F3-FZ |  |
| 19 F4-FZ |  |
| 20 F4-C4 |  |
| 21 F7-F3 | Frontal |
| 22 F8-F4 |  |

It should be noted that the EEG has a low spatial resolution. The grouping of the EEG bipolar channels in Table I is done according to the location of the two electrodes in the respective bipolar channel and the underlying cortical region of the brain. The cortical region is only used as a location marker for the EEG bipolar channel that measures the propagated electrical activity on the overlying scalp region. Therefore, in this study when results are presented with respect to the cortical region it does not refer to the explicit activity in the actual brain cerebral cortex.

### A. Kernel (dis)similarity matrices of HC and AD groups

Fig. 4 illustrates the kernel (dis)similarity matrices across all three epochs of the 20 HC and 20 AD participants used in this study. The kernel (dis)similarity matrices quantify synchrony between channels using spatiotemporal local similarities and global dissimilarities (Section III C). Thus, a measure of spatio-temporal functional connectivity.

**FIG. 4.**
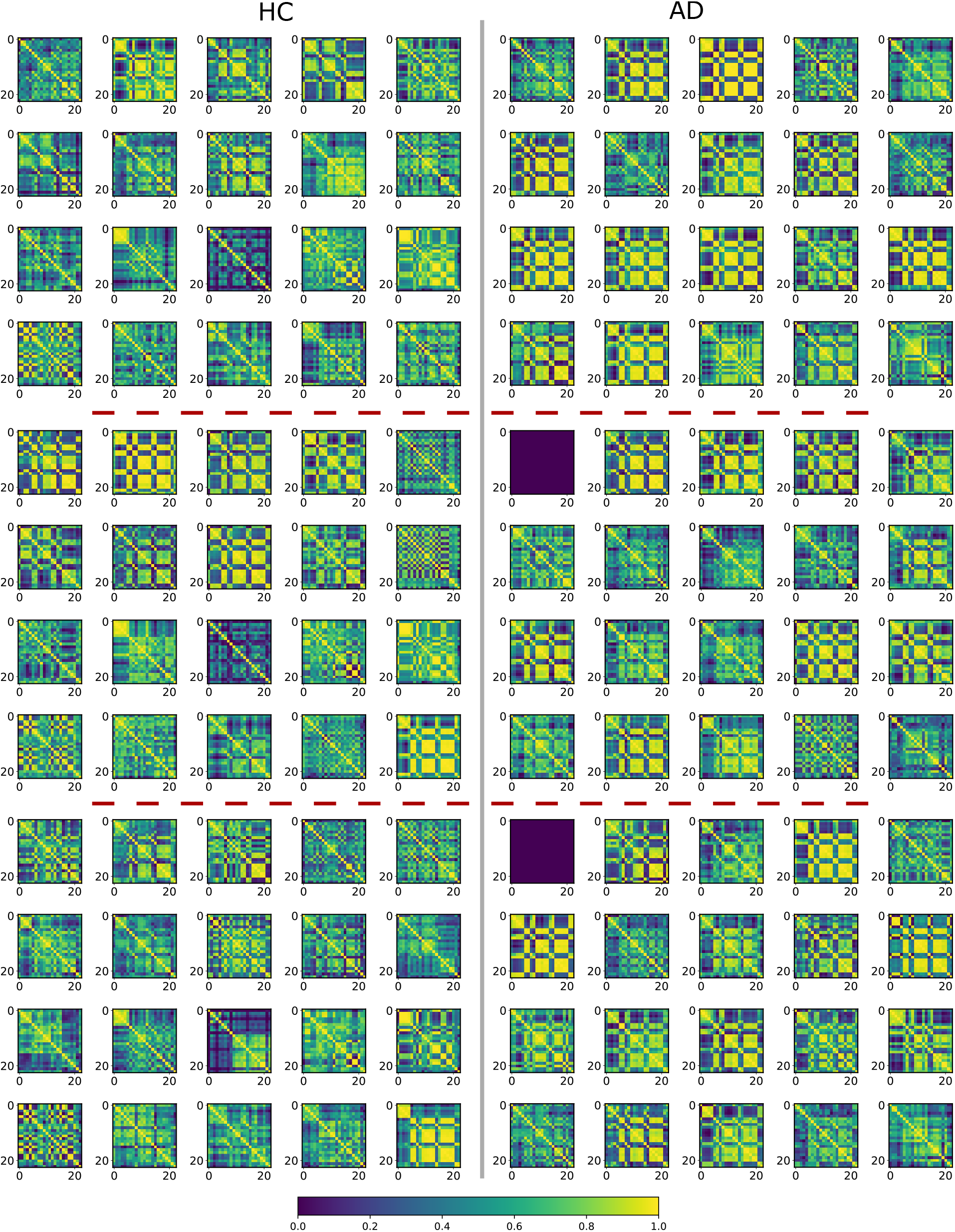
Participant specific normalised RBF kernel (dis)similarity matrices from Isomap-GPLVM of all epochs from all HC and AD participants for the best initial condition, eye-open data. All kernel (dis)similarity matrices are normalised such the diagonal is 1. The red dotted lines separate each set of respective epochs. It is evident that in a general sense the AD group does exhibit a certain pattern of spatio-temporal (dis)similarity or synchrony. This leads to the conclusion that this is a reflection of the specific patterns of dysfunction the literature mentions about AD patients – less dynamic complexity. The EEG data for epochs 2 and 3 of one participant in the AD group was not available. This is indicated by the all-zero kernel (dis)similarity matrices.

Statistically comparing the kernel (dis)similarity matrices between AD and HC groups, the significance-matrix **S** (Section III D) and the corrected significancematrix 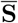 (after accounting for multiple comparisons, Section III D) is obtained. The Mann–Whitney U test was used to determine which elements of the kernel (dis)similarity matrices (pairwise comparisons–EEG channels) between HC and AD have significant differences. These significant elements are denoted as 1’s in **S**, zero otherwise. Fig. 5 illustrates **S**–blue indicates the significant elements. 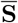 is obtained by using the BenjaminiHochberg false discovery rate controlling method (Section III D) to account for the multiple comparisons problem. Fig. 6 illustrates 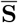–blue indicates the corrected significant elements.

**FIG. 5.**
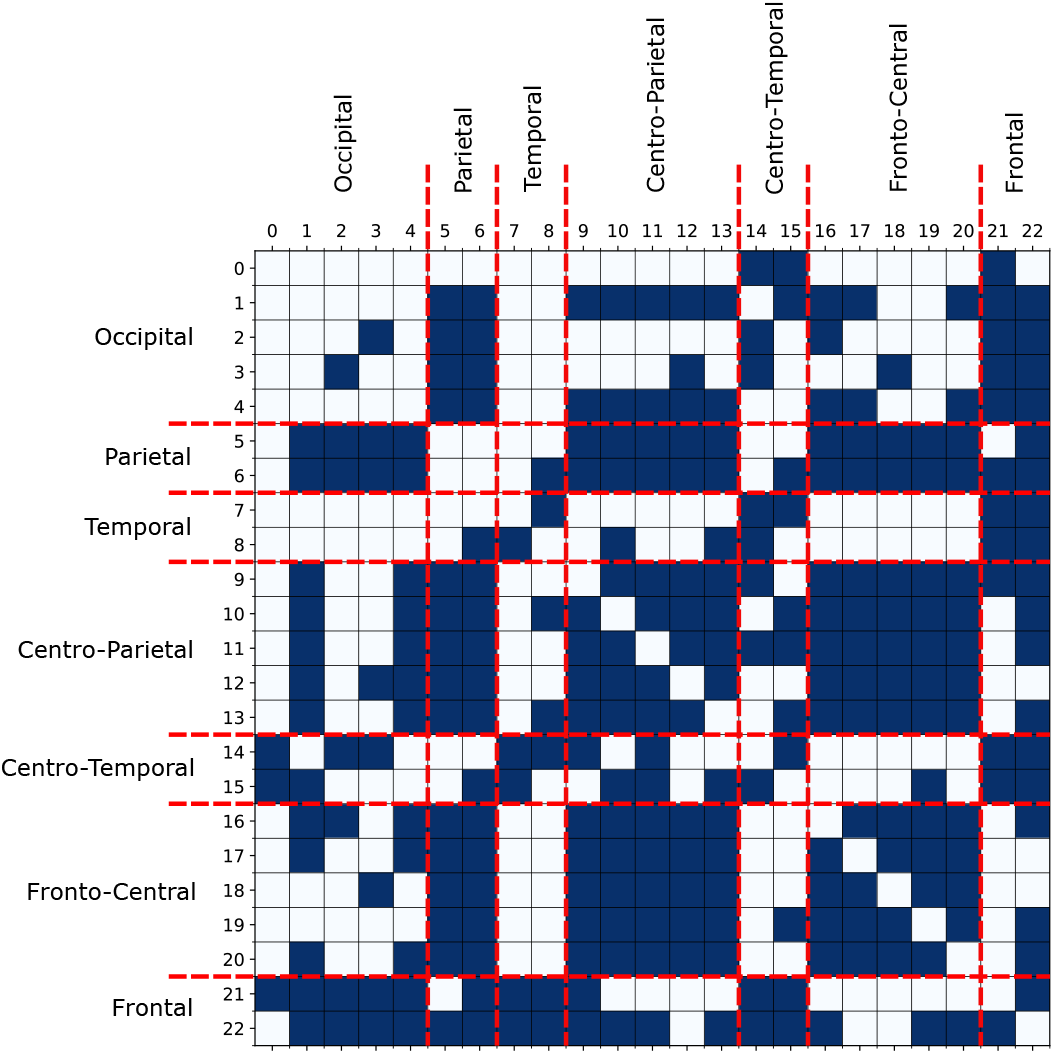
The significance matrix, S. This matrix shows the results of the element-wise statistical comparison (The Mann–Whitney U test, Section III D) of all the kernel (dis)similarity matrices (all epochs of all participants) from the HC and AD groups. The respective channel indexes are shown in Table I. The significant elements are denoted by 1 in **S**, illustrated in blue.

**FIG. 6.**
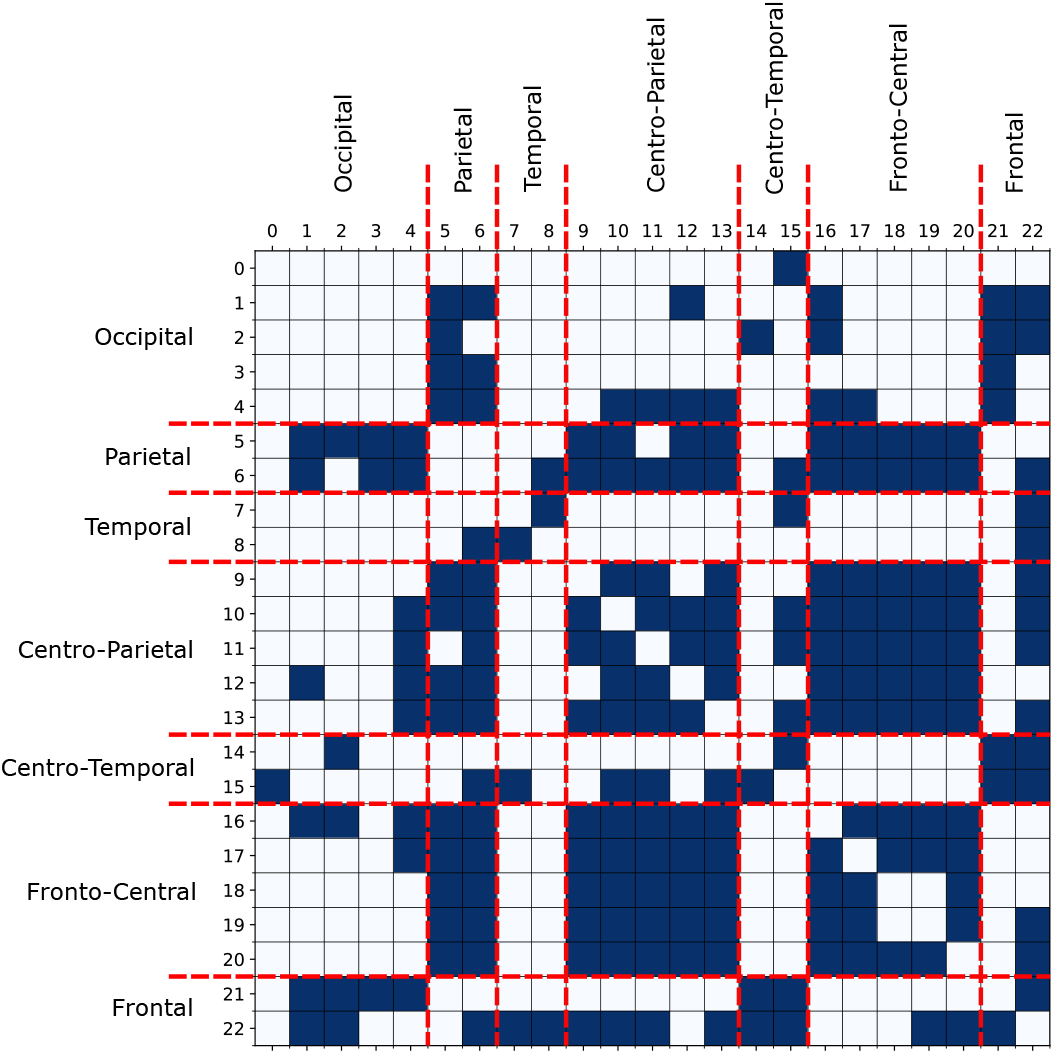
The corrected significance matrix, 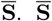. 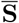 is obtained by applying the Benjamini-Hochberg false discovery rate controlling method due to multiple statistical comparisons. The respective channel indexes are shown in Table I. The corrected significant elements are denoted by 1 in 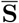, illustrated in blue.

Based on 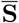 in Fig. 6, it is evident that there is a specific pattern in the statistically significant local and global variations in the functional connectivity between HC and AD. This can be a reflection of the specific patterns of dysfunction that have been mentioned in the literature, in which AD EEG data exhibits a specific change in synchrony compared to HC controls [2, 8, 25] and complexity in certain regions of the EEG being affected [18– 23]. These synchrony changes could be related to crossfrequency coupling [30, 33]. That, in turn, relates to local synchrony changes within and global synchrony changes between underlying cortical regions [30].

### B. Global functional connectivity changes and channel pair selections

The results in Section III F, in which kernel (dis)similarity values of a subset in channel pair combinations are used as features in linear SVM classification. This is to determine, in a global sense, which synchrony (spatio-temporal functional connectivity) differences between cortical regions (EEG channels) are more important in distinguishing between HC and AD groups. Fig. 8 illustrates the normalised averaged linear SVM weights (Section III H) of all the selected (using **S**) channel pairs in a matrix format. Table II shows the top 20 channel pairs that are ranked according to the normalised averaged linear SVM weights.

**TABLE II.**
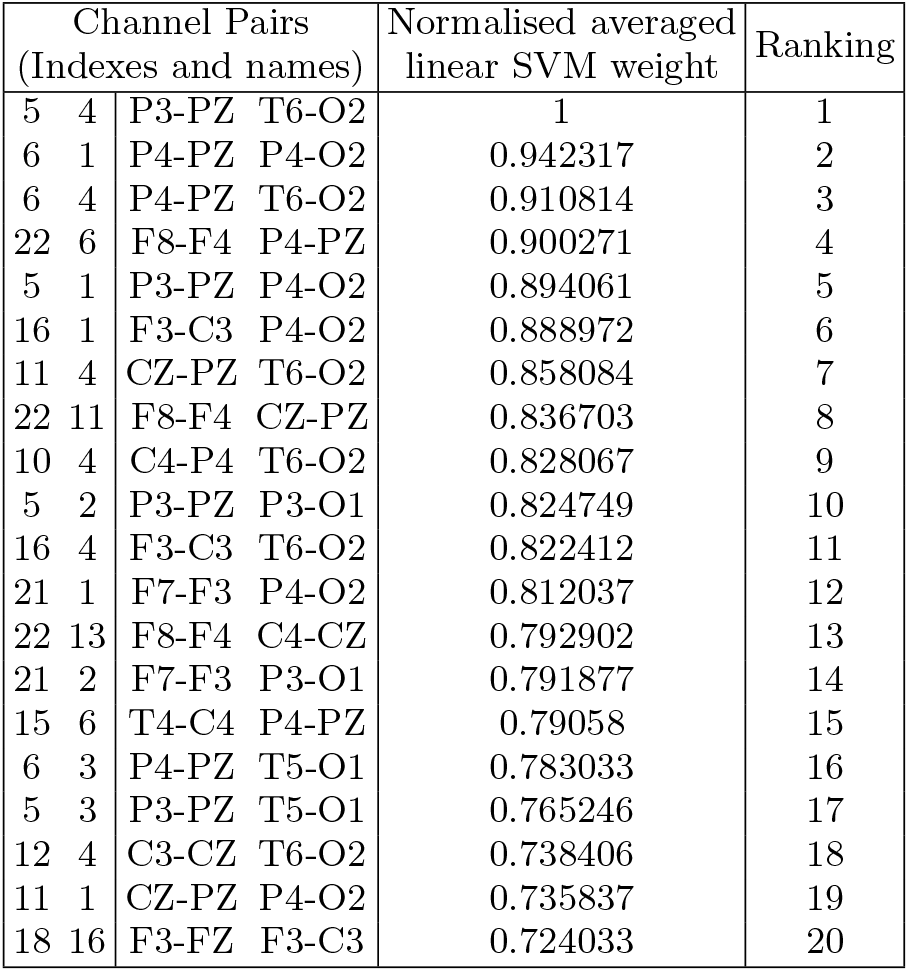
Ranking of (dis)similarity features of channel-pairs only the top 20 are shown

| Channel Pairs<br>(Indexes and names) |  |  |  | Normalised averaged<br>linear SVM weight | Ranking |
| --- | --- | --- | --- | --- | --- |
| 5 | 4 | P3-PZ | T6-O2 | 1 | 1 |
| 6 | 1 | P4-PZ | P4-O2 | 0.942317 | 2 |
| 6 | 4 | P4-PZ | T6-O2 | 0.910814 | 3 |
| 22 | 6 | F8-F4 | P4-PZ | 0.900271 | 4 |
| 5 | 1 | P3-PZ | P4-O2 | 0.894061 | 5 |
| 16 | 1 | F3-C3 | P4-O2 | 0.888972 | 6 |
| 11 | 4 | CZ-PZ | T6-O2 | 0.858084 | 7 |
| 22 | 11 | F8-F4 | CZ-PZ | 0.836703 | 8 |
| 10 | 4 | C4-P4 | T6-O2 | 0.828067 | 9 |
| 5 | 2 | P3-PZ | P3-O1 | 0.824749 | 10 |
| 16 | 4 | F3-C3 | T6-O2 | 0.822412 | 11 |
| 21 | 1 | F7-F3 | P4-O2 | 0.812037 | 12 |
| 22 | 13 | F8-F4 | C4-CZ | 0.792902 | 13 |
| 21 | 2 | F7-F3 | P3-O1 | 0.791877 | 14 |
| 15 | 6 | T4-C4 | P4-PZ | 0.79058 | 15 |
| 6 | 3 | P4-PZ | T5-O1 | 0.783033 | 16 |
| 5 | 3 | P3-PZ | T5-O1 | 0.765246 | 17 |
| 12 | 4 | C3-CZ | T6-O2 | 0.738406 | 18 |
| 11 | 1 | CZ-PZ | P4-O2 | 0.735837 | 19 |
| 18 | 16 | F3-FZ | F3-C3 | 0.724033 | 20 |

After evaluating different settings of linear SVM and Monte-Carlo cross-validation (Section III H) as described in Section III F, the selection of a subset in channel pairs using **S** along with an L2 penalty results in the best AUROC, 0.726. Subset selection avoids the channel pairs that are not statistically significant in distinguishing between HC and AD groups. However, the selection of **S** over 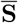 may be due to the Benjamini-Hochberg method being conservative [41], and could miss some actually important channel pairs.

According to the literature, in the case of AD, while changes in synchrony between EEG channels occur globally [8], single channel complexities, such as measures of entropy, mutual information and amplitude couplings, occur in channels relating to certain underlying cortical regions [24]. Some of these single channel complexity changes are:

- Increased amplitude coupling between delta and gamma bands across temporal, parietal, and occipital regions [18].
- Decreased amplitude coupling within the alpha band in the parietal region [22].
- Reduced auto-mutual information in channels within temporal, parietal, and occipital regions (referential montage channels T5, T6, O1, O2, P3, P4) [21].
- Reduced approximate entropy in channels within parietal, and occipital regions (referential montage channels O1, O2, P3, P4) [21].
- Reduced permutation entropy in temporal, parietal, and occipital regions [19, 20].
- Reduced sample entropy in channels within parietal and occipital regions (referential montage channels O1, O2, P3, P4) [46].

Therefore, in summary, single channel complexity changes occur mainly in the EEG channels relating to the temporal, parietal, and occipital regions. The EEG electrodes relating to these regions are shown in Fig. 7, highlighted in black–T5, T6, O1, O2, P3, P4. It should be noted the above single channel complexity studies were carried out using EEG channels from a referential montage. Interestingly, a prominent feature observed in Fig. 8 is the weighting of the functional connectivity changes between the parietal and occipital bipolar channels, Fig. 7–orange and purple lines respectively. The respective pairwise comparisons, between these bipolar channels, are ranked within the top 10 in Table II. Furthermore, these bipolar channels are spatially closely located along the scalp surface (see Table I and Fig. 3). As seen from Fig. 7, these bipolar channels are formed from the aforementioned referential EEG montage channels that, according to the literature, have reduced complexity on single channel analysis. From this study, it is observed, that the synchrony between the Parietal region (Fig. 7–orange) and the Occipital region (Fig. 7–purple) has significant changes between HC and AD. Specifically inter-regional synchrony (functional connectivity).

**FIG. 7.**
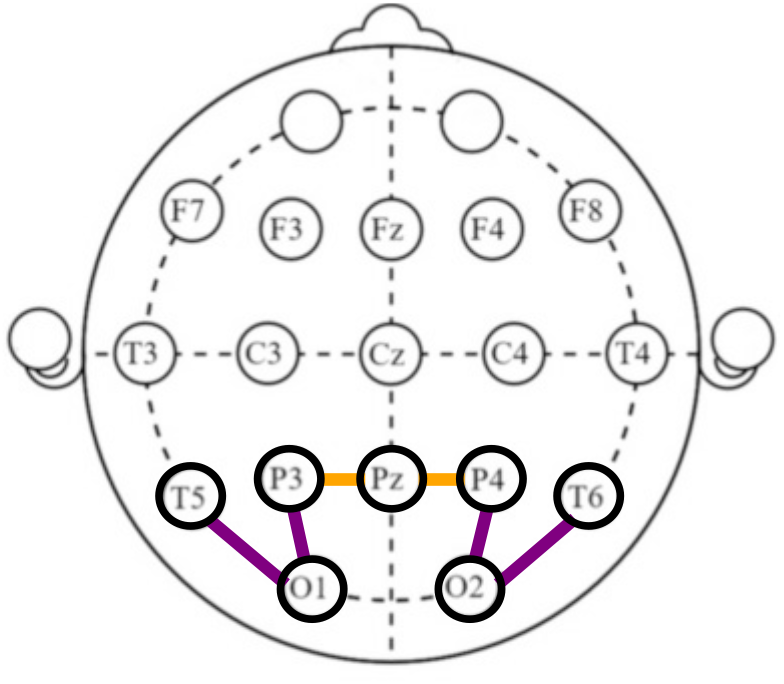
EEG electrodes and bipolar channels in the Occipital, Parietal and temporal regions. This illustrates the EEG electrodes, highlighted in black, relating to the EEG single channel complexity reduction in AD–according to literature. The orange and purple lines, connecting two nodes, indicate the bipolar channels respectively within the Parietal and Occipital regions (Table I) that also appear in Table II

**FIG. 8.**
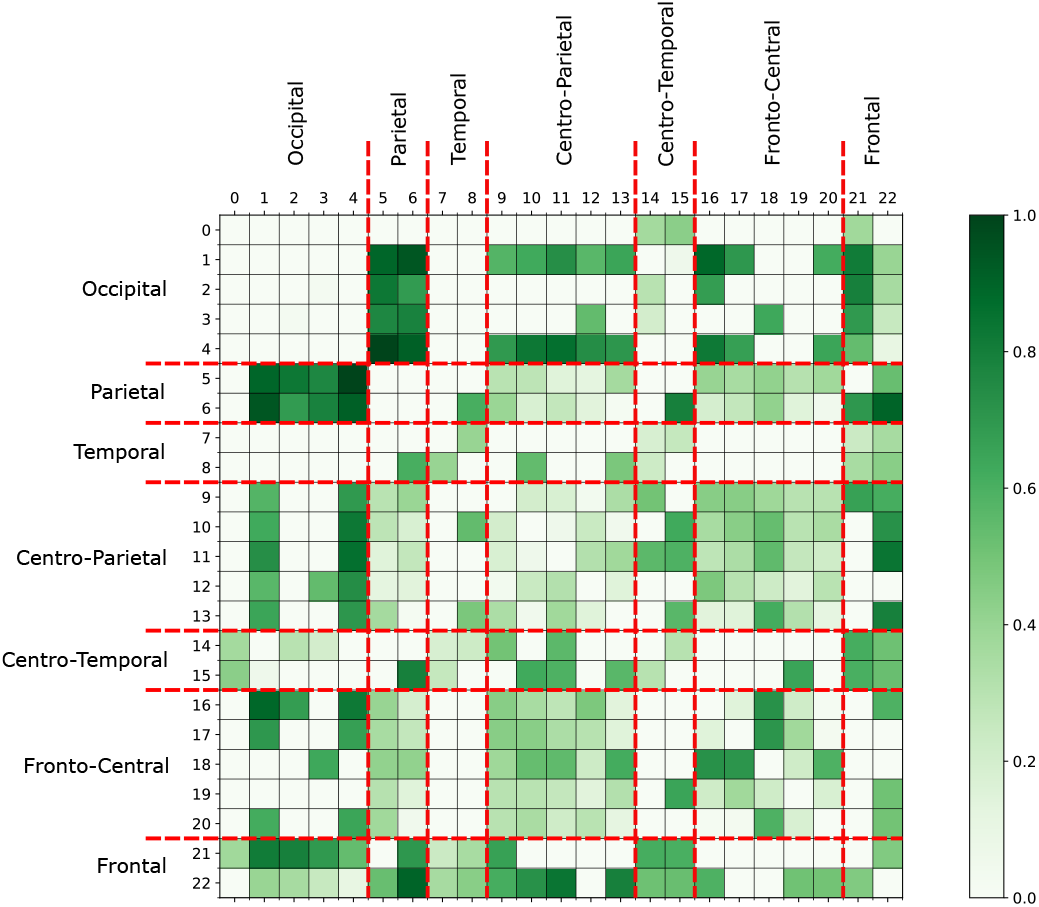
The matrix of normalised averaged linear SVM weights. This illustrates the weighting given by linear SVM to the kernel (dis)similarity features of the subset of channel pairs (Section III F). A prominent observation is the weighting on the Parietal and Occipital channel pairs.

Through observing the significance matrix, **S** (Fig. 5), and the corrected 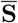 (Fig. 6), there are significant synchrony changes in AD within the Frontal and Occipital regions and between Frontal and Centro-Parietal regions. These channels are spatially widely distributed along the scalp surface and appear among the significantly weighted channel pairs, as in Table II. Therefore, the findings from this study reveal, at an EEG sensor level, clear changes in synchrony between long range cortical regions and also locally within adjacent regions. This matches the findings from the aforementioned literature. The importance of the significant synchrony (i.e. spatio-temporal functional connectivity) changes observed from **S**, Fig. 5, can be determined from the normalised averaged linear SVM weights (Section III H), which are reported in Fig. 8 and Table II.

### C. Channel-specific functional connectivity changes and channel pair selections

In this section, the linear SVM results by using the channel-specific approach (Section III G) to the analysis of functional connectivity from kernel (dis)similarity matrices are presented. This approach determines significant region-specific synchrony changes between HC and AD groups. Specifically, identifying which particular cortical regions (EEG channels), with respect to their interrelationships with other regions, can distinguish well between HC and AD groups. The channel-specific approach provides another layer of information, the other side of the same coin.

The rows of the kernel (dis)similarity matrices are used to form channel-specific feature spaces (Section III G) to be assessed individually via linear SVM and Monte-Carlo cross-validation (Section III H). This enables to identify which individual EEG channels have distinguishable changes in synchrony with other channels between HC and AD groups–the AU-ROC of the feature space. The normalised average linear SVM weights (Section III H) of the channel-specific feature space give a ranking of importance (in distinguishing between groups) for the inter-relationships with respect to that channel.

The AU-ROC of the channel-specific feature spaces is reported and used for comparison, as shown in Table III. Fig. 10 illustrates the most important channel inter-relationships in the top performing feature spaces, AU-ROC *>* 0.7 (highlighted in grey, Table III). These feature spaces correspond to the bipolar channels along the fronto-parietal regions, which are shown in Fig 9.

**TABLE III.** The average AU-ROC values of the channel-specific feature spaces

| Channel-specific feature space |  | Average AU-ROC of testing set |
| --- | --- | --- |
| 0 | O1-O2 | 0.604317 |
| 1 | P4-O2 | 0.650388 |
| 2 | P3-O1 | 0.635842 |
| 3 | T5-O1 | 0.615617 |
| 4 | T6-O2 | 0.612996 |
| 5 | P3-PZ | 0.652343 |
| 6 | P4-PZ | 0.6389 |
| 7 | T3-T5 | 0.581199 |
| 8 | T4-T6 | 0.646965 |
| **9** | **C3-P3** | **0.755004** |
| **10** | **C4-P4** | **0.734946** |
| **11** | **CZ-PZ** | **0.74525** |
| 12 | C3-CZ | 0.68545 |
| 13 | C4-CZ | 0.67407 |
| 14 | T3-C3 | 0.666313 |
| 15 | T4-C4 | 0.682305 |
| **16** | **F3-C3** | **0.72964** |
| **17** | **FZ-CZ** | **0.72943** |
| 18 | F3-FZ | 0.670291 |
| 19 | F4-FZ | 0.631998 |
| **20** | **F4-C4** | **0.735953** |
| 21 | F7-F3 | 0.616483 |
| 22 | F8-F4 | 0.660214 |

**FIG. 9.**
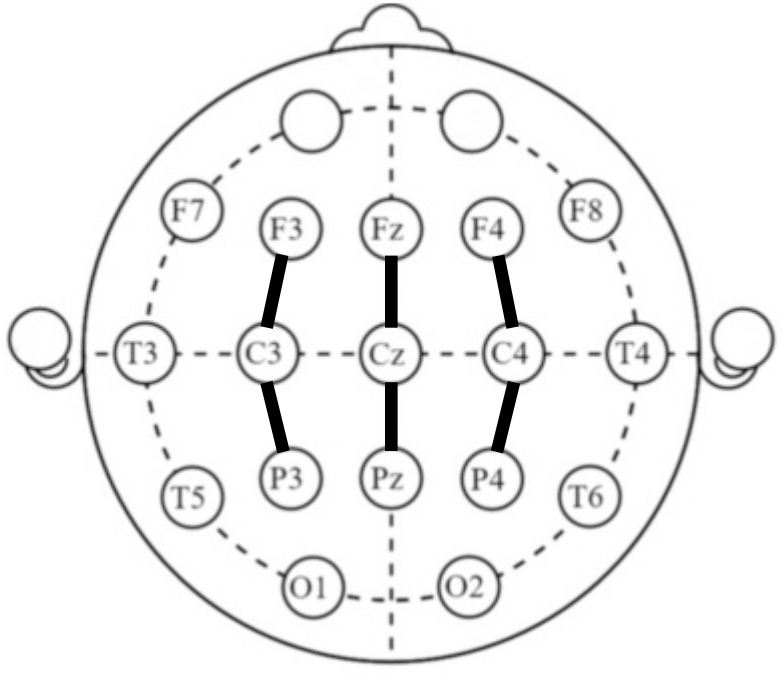
Underlying cortical regions of the top performing channel-specific feature spaces. The bipolar channels of the best performing channel feature spaces are mapped into the 10-20 international electrode placement. These channels lie within the fronto-parietal regions of the cortex.

**FIG. 10.**
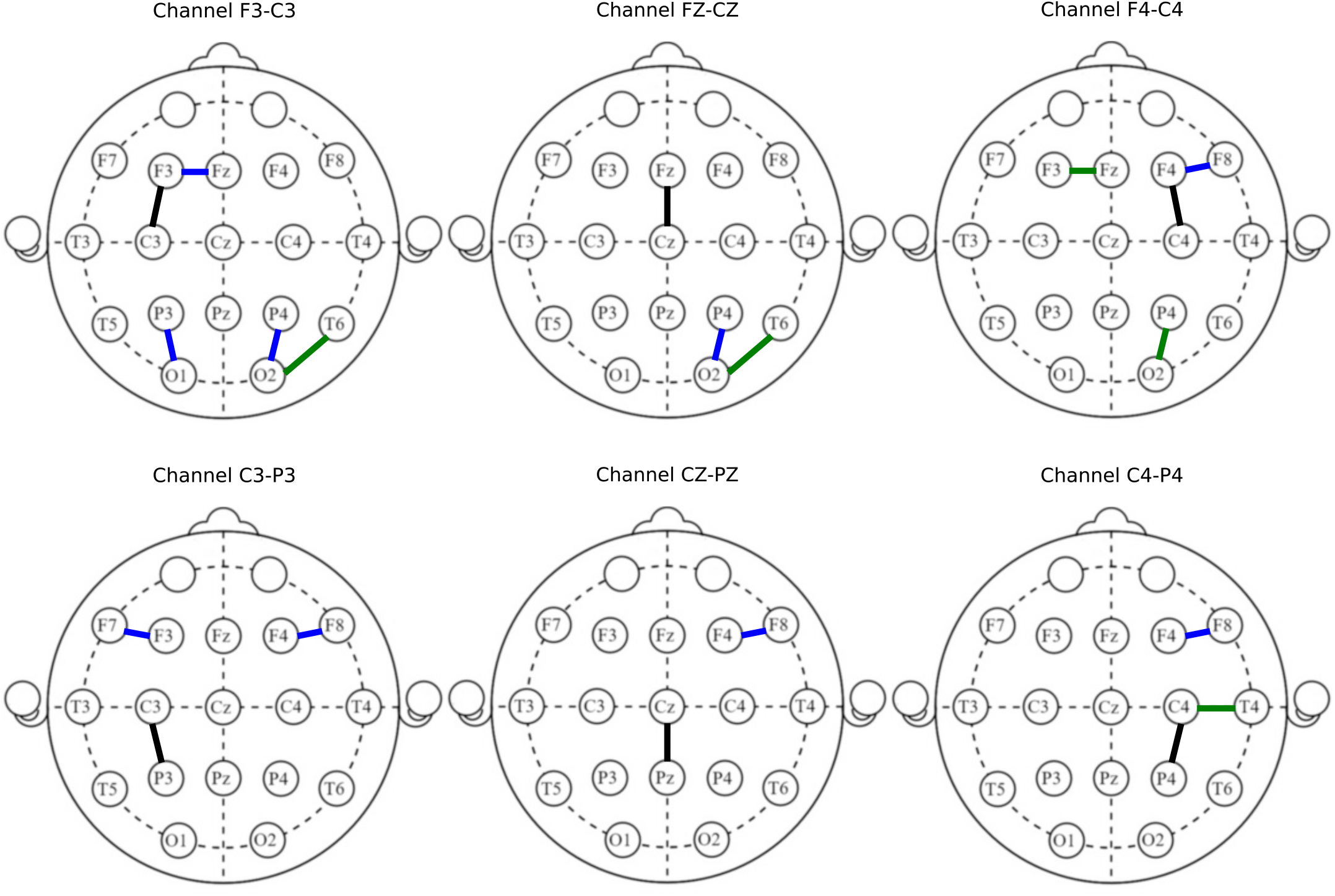
Important channel inter-relationships in the top performing channel-specific feature spaces. The channel inter-relationships with the highest weightings in the respective channel-specific feature spaces, AU-ROC *>* 0.7 are illustrated. These feature spaces (indicated in black) are from the channels within the fronto-parietal region. Black indicates the respective channel-specific feature space being considered. Channel pairs with, respect to the black channel, that has a normalised averaged linear SVM weighting of 1-0.9 are indicated by blue and 0.9-0.7 are indicated by green.

The fronto-parietal network has been reported to play an important role in the early diagnosis of AD in several studies using functional magnetic resonance imaging (fMRI) [42] and EEG [43, 44]. Neufang *et al*. [42] pointed out, at early stages of AD, regional gray matter volume is related to the reduction in effective connectivity (via dynamic causal modelling) in fronto-parietal networks. While Babiloni *et al*. [44] found that a measure of nonlinear inter-dependence (via the synchronisation likelihood) significantly reduces within the fronto-parietal channels of eyes-closed EEG data in AD patients.

In this study, we observed that the functional connectivity (i.e. kernel (dis)similarity matrices via manifold learning) changes between the channels in the frontoparietal region (Fig. 9) and other regions have a significant ability to distinguish between AD and HC groups in the eyes-open EEG. The functional connectivity changes with channels in the fronto-parietal region that have the highest normalised averaged linear SVM weights are illustrated in Fig 10. The bipolar channels in the centroparietal part of the fronto-parietal (i.e. C3-P3, CZPZ and C4-P4) have higher weights on the synchrony changes with frontal bipolar channels. While the frontocentral part of the fronto-parietal channels (i.e. F3-C3, FZ-CZ and F4-C4) have higher weights on the synchrony changes with bipolar channels from the occipital region. These are spatially widely distributed channels in the scalp. This is with the exception of channels F3-C3, F4C4 and C4-P4, which have higher weights on synchrony changes between channels that are spatially close as well.

## V. Conclusion and Future Works

A novel EEG channel selection methodology using kernel-based nonlinear manifold learning is presented in this work. This channel selection method aims to determine which spatio-temporal channel inter-relationships are better at distinguishing between HC from patients with AD for further dynamical analysis (e.g. causality, frequency coupling, nonlinear frequency response analysis). It was shown how a kernel-based spatio-temporal (dis)similarity matrix via manifold learning can be used as a measure of spatio-temporal synchrony (functional connectivity) between EEG channels, and can be used to determine which inter-relationships are important to characterise patients with AD.

The methodology presented can determine the changes in cortical (EEG channel) inter-relationships that are important in distinguishing AD patients from HCs. Also, it can be used to discover which specific cortical regions, at an EEG sensor level, can distinguish well between AD and HC groups. Furthermore, the results reported in our study are consistent with other studies in the literature. A main advantage of the proposed method is that the channel selection process takes account of both global and local spatio-temporal structures within the EEG data, to determine which linear and nonlinear channel inter-relationships are significant.

Through statistical comparisons of the kernel (dis)similarity matrices between groups, it is evident that the AD group exhibits a certain pattern of changes in synchrony. It was discovered that considering, bipolar derivations commonly used in clinical practice, functional connectivities between bipolar channels of temporal, parietal and occipital regions can distinguish well between AD and HC groups. Furthermore, the synchrony between bipolar channels of the fronto-parietal region and the rest of the cortical surface are important in diagnosing AD.

The main purpose of this paper is to introduce this novel channel selection methodology and its computational procedure. In-depth neuro-physiological interpretations with respect to AD were not discussed in the current study and this will be part of future work. Results of only EO data from three epochs for each participant were examined. Examination of EC data and a comparison with EO along with comparisons of different time windows (within the 12-second epochs) will be investigated in future work. Another important future study is to further investigate the detailed forms of nonlinearity using nonlinear dynamic modelling [47] and causality measures in both time and frequency domains [4, 48, 49]. These in-depth dynamical analysis methods will be applied to the channel pairs and regions determined using the Isomap-GPLVM method. Thus, enabling the further study of the underlining dynamic processes and connectivity in patients with Alzheimer’s disease. This will aid in identifying which linear and nonlinear dynamic features can be used in the diagnosis of the prodromal stages of AD.

## ACKNOWLEDGEMENTS

RG and FH acknowledge Coventry University for the Trailblazer PhD studentship. The EEG data was funded by a grant from the Alzheimer’s Research UK, grant reference number ARUK-PPG20114B-25. This is a summary of independent research carried out at the NIHR Sheffield Biomedical Research Centre (Translational Neuroscience). The views expressed are those of the author(s) and not necessarily those of the NHS, the NIHR or the Department of Health.

